# Mitigating Bias in Spatial Transcriptomic Pipelines via Human Feedback

**DOI:** 10.64898/2026.01.15.699786

**Authors:** Pierre Boyeau, Stephen Bates, Can Ergen, Michael I. Jordan, Nir Yosef

**Affiliations:** Department of EECS, University of California, Berkeley; Eric and Wendy Schmidt Center, Broad Institute, Cambridge; Department of Statistics, University of California, Berkeley; Department of EECS, Massachusetts Institute of Technology, Cambridge; Center for Computational Biology, University of California, Berkeley; Inria, Paris; Department of Systems Immunology, Weizmann Institute of Science, Rehovot

## Abstract

Biological discovery from experimental data, particularly large-scale assays, requires extensive preprocessing, during which raw outputs (e.g., images, sequences) are processed into structured forms that are more amenable to analysis. While statistical methods for such processed data are at the core of computational biology, the problem of coping with uncertainties introduced during preprocessing is a significant and underexplored issue. We address this issue in the context of differential expression analysis in spatial transcriptomics, which depends on a series of preprocessing steps, including demarcation of cell regions (segmentation), quantification of gene expression in cells, and cell-type annotation. We introduce Corrected Spatial Differential Expression (CSDE), a method that builds on Prediction-Powered Inference to leverage a small set of expert-validated data points (cells) to account for uncertainty due to preprocessing errors. Using two case studies, we demonstrate that CSDE produces more reliable and calibrated estimates of differential expression compared to the prevalent approach that neglects the impact of preprocessing. CSDE incorporates an efficient workflow to generate the required expert-annotated data, and is available as open-source at https://github.com/YosefLab/CSDE.

## 1 Introduction

Almost any assay in experimental biology requires a certain degree of processing to convert its raw outputs into a form that can be more easily perceived, analyzed, and visualized. A prevalent example of this is sequencing-based assays, such as RNA sequencing, whose raw output consists of strings of DNA that are converted through a series of steps into a quantification of gene expression. The gaps between raw RNA-seq data and its processed form (count matrices) have indeed been closely inspected [1, 2], and estimation of effects on downstream analyses has motivated efforts for uniform re-processing of RNA-seq data in the public domain [3].

Similar to RNA sequencing, fluorescent-based spatial transcriptomics (ST) also requires a series of preprocessing steps [4, 5, 6, 7]. Starting from raw images, many types of analysis necessitate a transformation of the data into a tabular form with estimation of expression (observed counts) of each gene in every cell. One major component of such a preprocessing pipeline is segmentation, namely, defining the boundaries of individual cells within the tissue. This process effectively assigns transcripts to their source cells, creating the distinct units whose gene expression will be compared. Algorithms for segmentation make use of cell morphological cues (stains for nuclei or membranes) and the captured transcripts (looking for recurring patterns of expression) [8, 9]. Once cells are defined, a subsequent step is to add labels of type or state, using their gene expression profiles [10, 11], which can be done manually or through automated tools [12]. The resulting annotated expression matrix can then be investigated, e.g., to study how gene expression in a given cell type differs between tissue regions [13, 14]. Errors during segmentation or annotation, however, can have substantial effects on the accuracy and validity of such interpretations [9, 15].

Several strategies have been considered to address these effects. One prevalent strategy takes a segmentation-free approach, operating directly on the captured transcripts and comparing tissue regions rather than individual cells [16, 17]. Another strategy attempts to improve the accuracy of segmentation, e.g., by increasing the complexity and scale of algorithms [18], datasets [19], features [8, 9], or supervision [20]. These improved methods, however, can and do still produce errors, e.g., due to high cellular density, overlap in the third dimension, irregularity of cell shapes, diffusion, and other forms of measurement noise. Recently, post-segmentation tools have emerged to help address this, aiming to identify and correct misassignments of transcripts to cells based on assumptions of regularity of expression in cell subsets [21] or subcellular distribution patterns [15].

Here, we present an alternative strategy for solving the problem: instead of attempting to correct the segmentation, we estimate the uncertainty it adds to downstream analysis, with a focus on differential expression. This method, CSDE (Corrected Spatial Differential Expression), takes a conservative approach, as it does not argue what the correct data should look like, but instead how incorrect our processed data may be. It has the advantage of flexibility, as it can account for any effects of preprocessing (e.g., annotation), rather than being restricted to segmentation alone. CSDE is motivated by and largely based on prediction-powered inference (PPI)—a recent framework for statistically valid inference from machine learning predictions [22, 23, 24]. PPI combines a small set of samples that were preprocessed by an expert (e.g., to determine their label) along with a much larger set of samples that were only preprocessed by an algorithm (e.g., predicting labels). The two sets of samples are used as input for some task (e.g., to estimate label-specific mean or quantiles), and the discrepancies between the two parts of the data are used to estimate uncertainty and to correct any biases that may have been introduced by the automated part, while still retaining statistical power.

CSDE builds on PPI to account for errors and bias in differential expression that result from inaccuracies at the two major preprocessing steps: segmentation and annotation. We implemented an interface for manual inspection of the data that provides a record of a small set of cells (samples) whose segmentation and assigned labels are deemed valid. We combined this with a much larger set of cells in the same dataset that were segmented and labeled by algorithms. Using lung and colon tumors assayed with the MERSCOPE platform, we demonstrate that CSDE can estimate differences in gene expression between regions with a higher degree of accuracy and reproducibility and without compromising power.

## 2 Methods

Our objective is to obtain statistically valid estimates of differential expression between spatially resolved cell subsets, specifically accounting for quantification errors introduced by automated analysis pipelines. Our framework, CSDE, illustrated in Figure 1, leverages two complementary data modalities: a large dataset of cell-by-gene expression quantifications that was obtained via an automated pipeline (providing statistical power but potentially suffering from systematic errors), and a small dataset obtained via manual curation (providing accuracy but limited in scale). Our analysis begins with a fluorescence-based spatial transcriptomics assay of one or several tissue samples.

**Figure 1:**
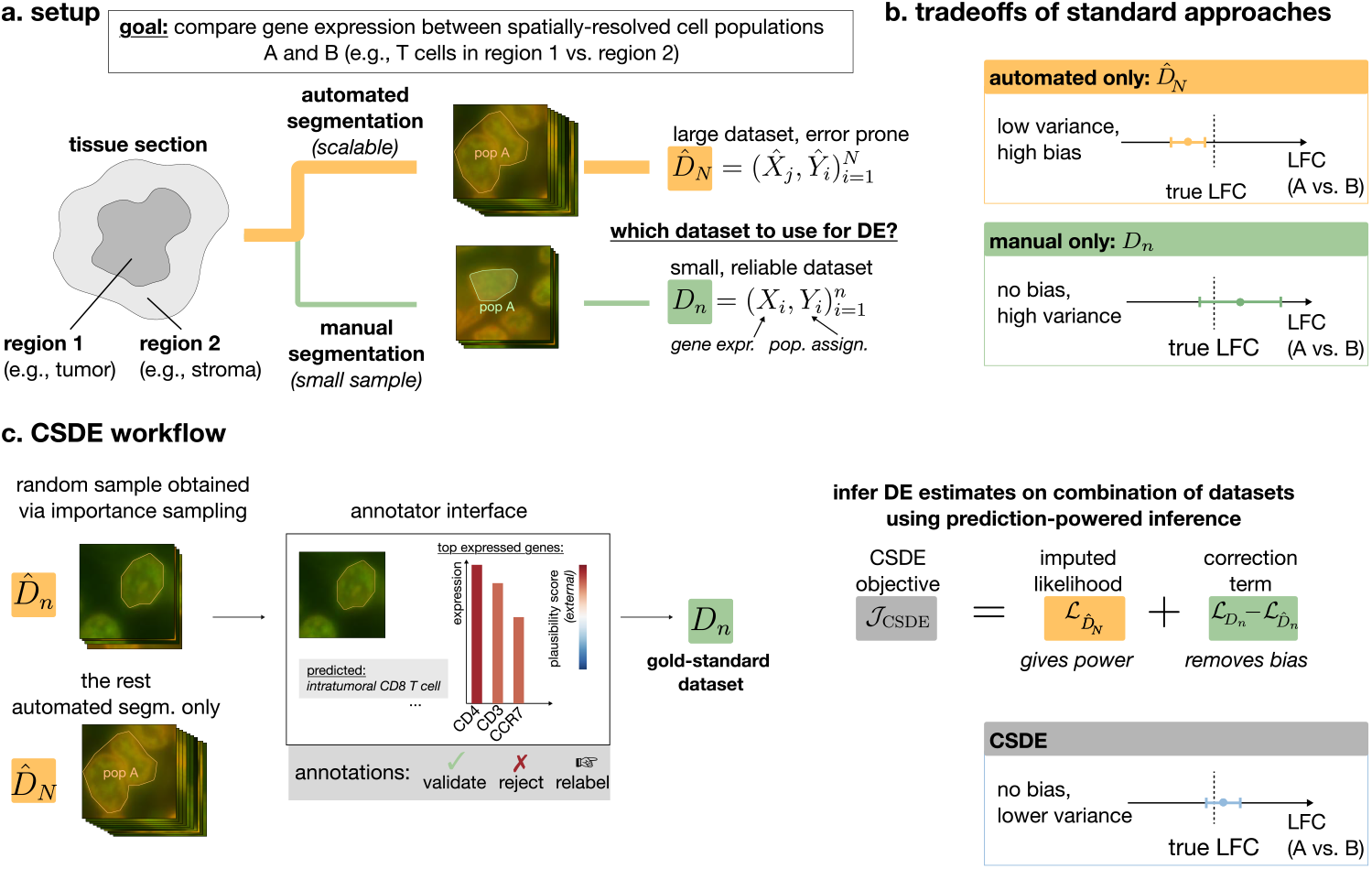
The CSDE model. **a**. We focus on the problem of differential expression (DE) between spatially resolved cell subsets from a spatial transcriptomics assay. Two approaches for cell segmentation are commonly used to obtain candidate cells: automated pipelines (high-throughput but error-prone) or manual segmentation (high-quality but low-throughput). **b**. The canonical parameters of interest for DE are log-fold changes (LFCs) of expression between subsets. LFCs quantify the magnitude of DE, where positive values indicate higher expression in the subset of interest, and negative values indicate lower expression. Automated approaches provide large sample sizes, yielding low-variance but potentially biased LFC estimates due to classification errors. Conversely, manual curation produces unbiased estimates but suffers from high variance due to limited throughput. **c**. CSDE combines the strengths of both approaches by first streamlining manual data curation. For this purpose, a small set of cell images is sampled via importance sampling, giving priority to cells more likely to contain the subsets of interest. For each image, a human expert validates, rejects, or corrects the automated results by visually inspecting the segmentation overlaid on the raw data, along with the predicted cell label and expression of top DE genes. CSDE then estimates LFCs by optimizing an objective that leverages the large automated dataset while removing bias using the manual labels. The resulting LFC estimates are both low-variance and consistent.

We assume the assay generates a collection of *candidate* data points 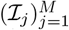, each centered on a *potential* location of a cell. These localizations are typically determined by an initial segmentation step and provide an initial guess of where cells are located. ℐ_*j*_ contains all the *raw information* around its central point, including location information and all fluorescence readouts (providing morphological cues and locations and numbers of transcripts). In the following, we will show how to use the bulk of the collection of data points to form our large and automated modality, and then how to inspect selected parts of the collection (using importance sampling) to generate the curated modality.

### Generating a large sample set through automated analysis

Establishing the automated (large) data modality relies on a pipeline with two (potentially combined) major steps: (i) identifying (segmenting) cells and estimating gene expression levels within each cell, and (ii) labeling cells according to their subset. Here, we define a cell subset as a flexible concept that defines the groups to be compared in a DE analysis. A subset can, for instance, represent a specific cell type (e.g., neurons vs. glia), a spatial region (e.g., cortical layers), or a combination of both (e.g., excitatory neurons in Layer V). Such pipelines normally range from proprietary software of commercial platforms (e.g., MERSCOPE or nanostring) to custom-built workflows that rely on open-source tools.

Formally, let 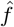 be the automated pipeline that produces gene expression quantification 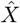 and a subset assignment *Ŷ* for every data point (candidate cell) in the collection such that^1^:

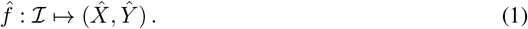

Here, ℐ denotes a single data point in the collection. 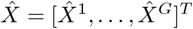 is a *G*-dimensional vector of gene expression estimates. In particular, 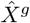 is the number of transcripts observed for the cell for gene *g* ∈ {1, …, *G*}. *Ŷ* ∈ {0, …, *K*} is the label assigned to the cell out of a vocabulary of *K* possible subsets plus an additional (*K* + 1-th) *technical* category (representing invalid or rejected cells) that we use during the execution of CSDE.

### Generating a small dataset with manual curation

As an automated pipeline, 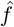 is susceptible to errors and systematic biases. Most crucially, inaccuracies in identifying cell boundaries (during cell segmentation) can lead to misassignment of transcripts and erroneous estimations of gene expression. During cell classification, errors can arise from two primary sources: the inherent imperfection of a classification model or a domain shift between the data used to train the annotation algorithm and the spatial transcriptomics data to which it is applied. These errors can often be identified by human experts via visual inspection of the raw data, which can be used to produce better-quality sets of cells, along with their gene expression profiles and assigned label. Formally, let *f* ^⋆^ be the manual procedure such that

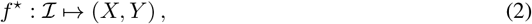

where *X* and *Y* are the manual gene expression and classification quantifications, respectively, following the same interpretation as in Equation (1). Unfortunately, manual curation is time-consuming and does not scale to the number of cells in typical ST experiments. However, as we demonstrate next, scrutinizing a small and randomly-sampled set of data points (i.e., *n* ≪*M*) from the collection within the CSDE framework is sufficient for effective analysis.

We partition the *M* data points into two sets. The first, the *manual set*, consists of the *n* data points 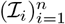 selected for manual curation. For these data points, both manual (*f* ^⋆^) and automated 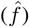 quantifications are available. The second, the *unlabeled set*, consists of the remaining *N* = *M* − *n* data points 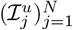 which are processed only by the automated pipeline 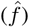. This partition gives rise to the three datasets used by CSDE:

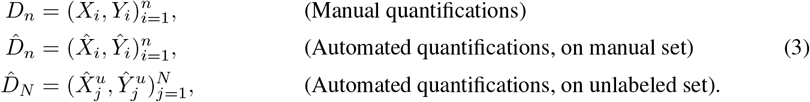

Here, we have (*X*_*i*_, *Y*_*i*_) := *f* ^⋆^(ℐ_*i*_) and 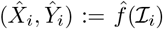 for any data point ℐ_*i*_ of the manual set, while 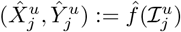 for any data point 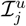 of the unlabeled set.

### 2.1 A scalable workflow for manual inspection of spatial transcriptomics

We now address the practical challenge of efficient manual inspection. Our approach is guided by the following principle: It is easier and more efficient for a human expert to judge the quality of an existing (automated) analysis (here, segmentation and labeling) against the raw data than it is to create a perfect one from scratch. We therefore designed the manual inspection pipeline in our case to act downstream and refine the output of an automated pipeline.

We devised a streamlined validation protocol designed to be both fast and reliable. To minimize expert time, we focus this step on a small and randomly-sampled subset of data points (putative cells). For each data point, the annotator takes one of three actions: If both the segmentation and the label appear correct, the annotator *accepts* the quantification, setting the manual data equal to the automated data 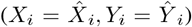. If the segmentation is adequate but the label is not, the annotator can *correct* the subset assignment, which updates the manual label while retaining the expression data 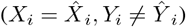. To keep annotation time low, our workflow does not support correcting segmentation boundaries. Finally, if a quantification is deemed unreliable, the annotator *rejects* it, and these cells are handled as a distinct category in our model (*Y*_*i*_ = *K*).

To make these decisions, the annotator reviews a composite diagnostic panel for each cell to be validated, which overlays the automated segmentation and classification results onto the raw image data. Figure 2 illustrates the panel used by annotators for validation on a dataset of human lung cancer profiled with MERFISH [25]. To illustrate the workflow, we consider the task of identifying genes that are differentially expressed by T cells in intratumoral vs. peritumoral regions. For the automated pipeline 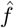, we used `proseg` [9] for segmentation, followed by a clustering-based assignment of labels (Supplementary Methods). In the example we present, the annotator is asked to validate an automatically processed T cell. To this end, the annotator first visualizes cell boundaries overlaid with cell membrane marker intensities (Figure 2a). If the segmentation is deemed adequate, the annotator validates the cell’s subset assignment by visualizing key markers in the surrounding tissue (Figure 2a) and exploring the gene expression profile of the center cell (Figure 2b).

**Figure 2:**
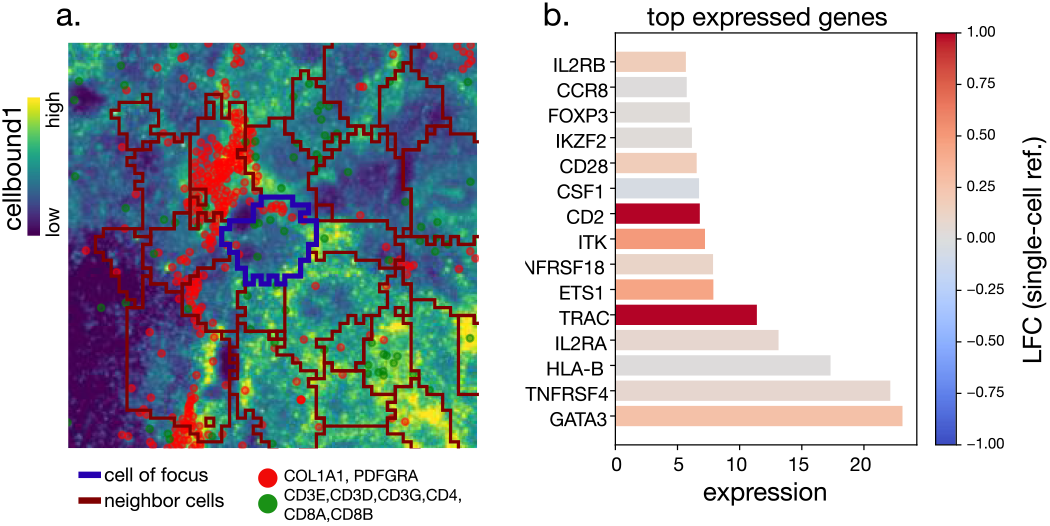
Example of images used for validation-based annotation.

Practically, we implemented this workflow by using a custom script to produce the diagnostic image panels for each cell to be validated. These image panels were uploaded to the Computer Vision Annotation Tool (CVAT) [26], a specialized platform for manual image annotation. After annotation, the labels were downloaded and integrated with the gene expression data for analysis.

### 2.2 Human-in-the-loop spatially-resolved differential expression

Our objective is to quantify differential expression (DE) between two putative cell subsets, denoted by the index sets *A, B* ⊆ {1, …, *M*} . To do so, CSDE employs a generalized linear model (GLM), a canonical approach for count-based DE:

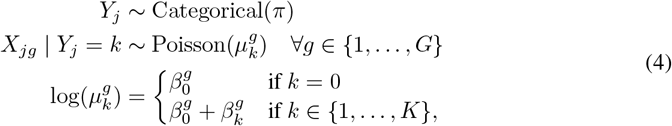

where *j, g*, and *k* are indices of cells, genes, and cell-subsets (resp.) and 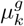 is the expected expression of *g* in cell subset *k*. Each cell is treated as an independent observation, and gene expressions are assumed to be independent conditional on the cell subset.

In this model, the parameter 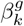 quantifies the log-fold change (LFC) in the expected expression of gene *g* for cells of subset *k >* 0 relative to the reference subset (*k* = 0). When comparing cell subsets *A* and *B, k* = 0 corresponds to subset *A, k* = 1 corresponds to subset *B*, and *k* = 2 to all other cells, including those rejected, e.g., due to segmentation errors, during annotation (see Section 2.1). The model therefore operates on all cells, not just those of primary interest (*A* and *B*), because a manual correction can re-assign a cell’s label to any other category.

Obtaining a point estimate and confidence interval for 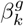 allows us to quantify the effect size (LFC) and test the null hypothesis 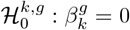, thereby calling DE genes. Traditionally, these quantities are estimated via maximum likelihood estimation (MLE) of any given gene *g* and dataset *D*:

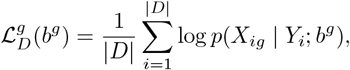

where *p*(*X*_*ig*_ | *Y*_*i*_; *b*^*g*^) is the gene expression probability and 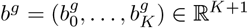 is the vector of LFC parameters.

The question that arises is: *which dataset of the three enumerated in Equation* (3) *should we use to estimate these parameters?* Using manual inspection (*D* := *D*_*n*_) leverages high-quality quantifications, but *D*_*n*_ might be too small to obtain useful (i.e., sufficiently tight) confidence intervals for 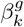, or, equivalently, to obtain a high statistical power to test for 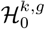. We have on the other hand access to a large dataset that has been processed automatically 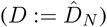. Because they are not reliable, however, these quantifications can introduce systematic bias in the estimation of 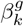. CSDE resolves this by optimizing a corrected objective function that integrates all available data using prediction-powered inference [22]:

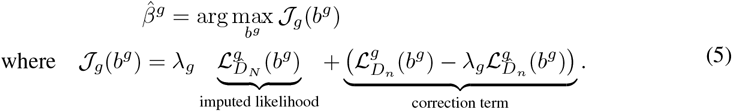

*λ*_*g*_ ∈ [0, 1] is a tunable hyperparameter that controls the importance of the automated dataset for parameter estimation, which we will discuss in more detail later in this section. The objective function 𝒥_*g*_(*b*^*g*^) that our estimator optimizes is a random variable that contains two terms. The first term is the imputed likelihood computed on the (unreliable) automated quantifications 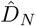. The second term is a correction term ensuring that the objective 𝒥_*g*_(*b*^*g*^) is not biased by the automated quantifications. Indeed, the CSDE objective function is on expectation equal to the log-likelihood of the model on the manual data:

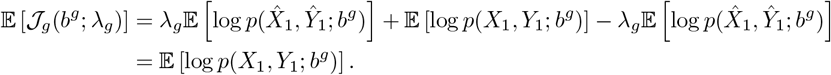

This property ensures that errors in the automated quantifications do not confound the analysis, as any bias they introduce is effectively canceled out in expectation. Beyond the unbiasedness of its objective function, CSDE benefits from useful statistical guarantees in the context of DE analysis. In particular, it provides consistent (meaning that the estimator converges to the true parameter value) and asymptotically normal (meaning we can build reliable confidence intervals or test hypotheses) estimates. The following corollary formalizes these guarantees:

#### Corollary 1

(Asymptotic normality of 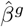, from [22]). *Suppose that the manual datasets D*_*n*_ *and the automated datasets* 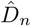 *and* 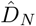 *are generated as above. Also assume that asymptotically, the ratio n/N converges to some constant r >* 0. *Then*, 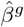 *from Equation* (5) *is asymptotically normal. In particular, for any given subset k and gene g, we have that:*

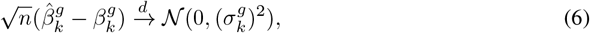

*where* 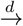 *denotes convergence in distribution and* 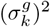 *is the asymptotic variance of* 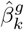 *that can easily be estimated from data*.

The asymptotic normality of 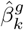 in Equation (6) provides a straightforward way to build confidence intervals, or equivalently, to test for 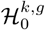.

The CSDE estimator thus combines the complementary strengths of the manual and the automated parts of the data. On the one hand, it provides consistent and asymptotically normal estimates, similar to a standard analysis of the manual data alone. Contrary to the standard approach, however, CSDE also leverages the large automated dataset to increase statistical power, providing confidence intervals that will be narrower on average than those obtained from the manual data alone [23].

### 2.3 Implementation details

The original approach [23] selects *λ* to minimize the total variance of the entire parameter vector, corresponding to the vector 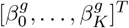 in our setting. In our setting, the parameter of interest is a single coefficient, 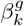, the log-fold change for cell subset *k* relative to the reference, rather than the entire parameter vector. Minimizing the total variance can result in over-conservative confidence intervals for this specific coefficient, potentially masking subtle differential expression signals. We therefore select *λ* to minimize the asymptotic variance of 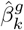, maximizing statistical power for the comparison of interest. More details on the selection of *λ*_*g*_ are provided in the Supplementary Information.

The construction of the manual dataset *D*_*n*_ is another critical implementation detail. An incorrect approach for creating the manual dataset *D*_*n*_ would be to sample only cells that are predicted to be in a subset of interest (*A* or *B*). The reason is that such pre-filtering will necessarily rely on the automated pipeline, which can assign the wrong labels to cells. For this reason, the manual dataset must be annotated based on a random sample of all cells. Uniform random sampling, however, can still be problematic if our subsets of interest cover a small portion of the cells in the tissue. To address this, we use importance sampling to select data points for manual annotation, giving priority to data points that are more likely to be of interest.

We designed a weighting function *w* allowing us to focus on images for which the automated annotation predicts cell subsets of interest. Concretely, the function assigns a weight *w*(ℐ) ∈ ℝ^+^ to each data point ℐ in the collection, such that the probability of selecting ℐ for manual annotation is proportional to *w*(ℐ). However, this non-uniform sampling scheme introduces a bias that must be corrected to ensure our estimates remain statistically valid. To do so, we adjust the objective function from Equation (5) by re-weighting each observation in the correction term using inverse weights:

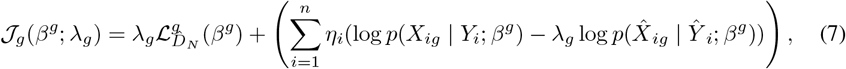

where 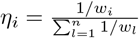 is the normalized weight of the *i*th manually annotated cell, with *w*_*i*_ = *w*(ℐ_*i*_).

A proof of the validity of this correction is provided in the Supplementary Information.

## 3 Results

To benchmark our approach, we focused on characterizing infiltrating CD8 T cells in lung and colon cancers, aiming to identify DE genes between intratumoral and extratumoral subsets. This task is particularly challenging due to the close proximity of T cells with antigen-presenting cells or tumor cells, interactions that are essential for their function. As documented elsewhere [8, 9], this crowded environment typically leads to segmentation and classification errors, introducing spurious DE signals that can mask true biological changes.

For this analysis, we extracted four slides (two for lung and two for colon) from the human immunooncology dataset of the MERSCOPE platform [25]. These slides provide information on the expression and location of a panel of 500 genes and include several additional cues of cell morphology through nuclear and membranal stains.

### Automated cell segmentation & classification

We used a combination of segmentation algorithms and canonical single-cell clustering and annotation methods to segment and classify cells. For segmentation, we employed Proseg [9], a state-of-the-art method that relies on transcript locations (rather than membrane or nuclear stains). After segmentation, we employed the Leiden clustering algorithm [27] on the log-transformed and median-normalized expression matrix, and then manually labeled them at coarse resolution using marker genes (Table S1). Notably, although the clusters were manually labeled, the obtained annotations are the direct product of an algorithmic approach to segment and cluster the cells. Indeed, difficulties in distinguishing cluster boundaries (e.g., distinguishing CD4+ from CD8+ T cells S1a) are a common problem due to which cells can be mislabeled during this step. To identify intratumoral and extratumoral regions, we stratified the tissue into tumor and adjacent regions based on the proximity of each cell to tumor cells identified by the automated pipeline (Supplementary Methods). These spatial assignments were treated as fixed and used to define cell subsets for all subsequent analyses, including manual curation.

### Manual annotation curation

For each slide, we built a *manual dataset* by validating, correcting, or invalidating the automated cell segmentation and type annotations for a random sample of 600 raw data points (cells) per slide. Since T cells were relatively rare (Table S2), we employed our importance sampling to prioritize cells identified as T cells by the automated pipeline. For each slide, we increased the weight of cells identified as T cells to ensure that approximately a third of the sampled cells were T cells (approximately 200 cells out of the 600 cells sampled in total). Using a widely-used image annotation tool as a user interface [26] on image panels like Figure 2, a single annotator evaluated each cell in the sample. First, the annotator determined if the segmentation was adequate. If so, they checked the cell’s label; if the automated label was incorrect, they assigned the correct one (whether T cell or otherwise). This process took less than an hour per slide.

### Benchmark

We compared CSDE with two baselines representing current approaches for spatial DE analysis. The first baseline, *automated*, identified DE effects by fitting the parameters of the GLM of Equation (4) on the automated dataset alone using maximum likelihood estimation. This method reflects the common practice of using automated segmentation and classification with minimal manual curation. The second baseline, *manual*, identified DE effects via maximum likelihood estimation using the same model, but using only the manual dataset. This approach represents a conservative approach, relying solely on high-confidence manual annotations (inherently limiting the number of cells analyzed). DE genes were identified for the two baselines and for the CSDE method as those with p-values below 0.05 after correction for multiple hypotheses [28].

#### 3.1 CSDE improves T cell characterization in lung cancer

We first focused on characterizing CD8+ T cell infiltration in lung cancer. The first slide served as our primary dataset for the main analyses, while the second was used as a replicate to assess reproducibility. After segmentation, the automated pipeline applied to the first slide returned a set of 351,309 data points (putative cells) that were coarsely annotated using clustering (Figures 3a, S1a). Manual analysis was done as above, for approximately 600 cells. Investigation of the manual annotations revealed significant segmentation and misclassification errors. Among the cells predicted as T cells by the automated pipeline and manually reviewed, only 47% were confirmed as correctly segmented and classified T cells. Cells misannotated as T cells by the automated pipeline included actual T cells with improper segmentation or classification, and non-T cells incorrectly classified as T cells (Figure S2).

**Figure 3:**
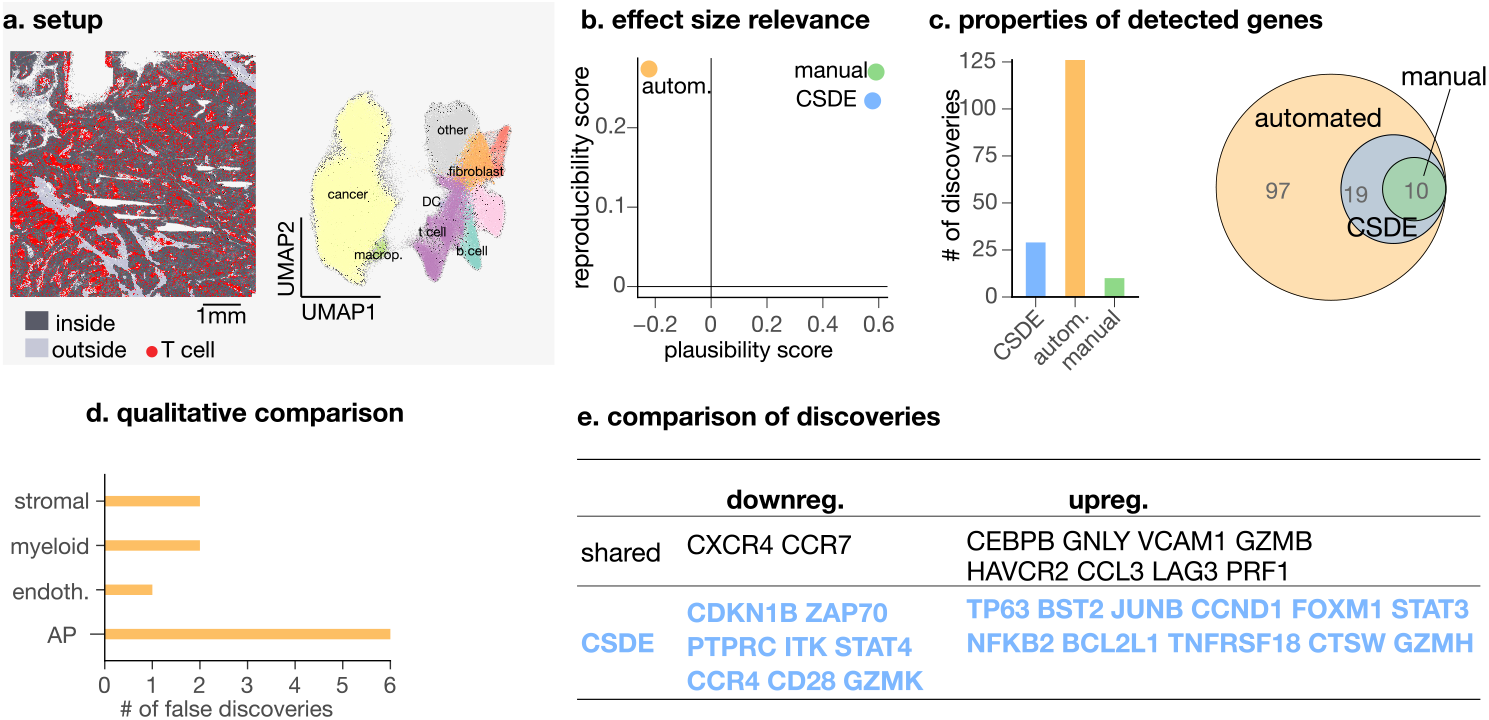
Lung experiment. Comparison of CD8 T cell gene expression profiles inside and outside tumors. **a**. *Left*: visualization of tumors in the tissue with CD8 T cells (obtained via automated pipeline) in red. *Right*: low-dimensional embeddings (UMAP) of the cells obtained by the automated pipeline. **b**. Relevance of the DE effect sizes obtained with the different methods based on reproducibiliy and plausibility scores. The biological plausibility score assesses the extent to which genes with large positive or negative LFCs are known to be expressed by T cells. The reproducibility score assesses the extent to which the LFC estimates are consistent between the two slides. See Supplementary Methods C.4 for more details. **c**. Properties of the discoveries made by the different methods based on their number (*Left*) and overlap (*Right*). **d**. Quantification of non-T cell marker hits for the discoveries made by the different methods. AP: Antigen-presenting. The list of non-T cell marker genes is provided in Table S3. **e**. Comparison of the discoveries made by our method and the *manual* approach for the comparison of CD8 T cells inside and outside tumors.

To set up the comparative analysis, we used the automated dataset to stratify the tissue into tumor vs. adjacent regions (Figure S1b; based on proximity to tumor cells; see Supplementary Methods). The *automated* benchmark detected the largest number of genes as DE, while the *manual* benchmark produced the lowest amount. This result is indeed expected given the difference in the number of cells that each method considered. CSDE identified fewer genes than the automated approach, but significantly more than the *manual* approach (29 versus 10 genes). Importantly, CSDE recovered all discoveries of the *manual* benchmark, suggesting that CSDE enhances detection power while maintaining biological validity.

To test this, we evaluated CSDE and its benchmarks using two scores. The first, the *plausibility* score, quantifies the extent to which genes with strong DE effects are known to be expressed by T cells, calculated as the Spearman correlation between estimated absolute log-fold changes and gene expression in T cells from a reference single-cell RNA-seq pancancer dataset [29] (Supplementary Methods). We introduce this score since transcript misassignment (e.g., assigning to T cells transcripts of genes that are not expressed by T cells) is a prevalent problem in spatial transcriptomics. The second score estimates *reproducibility* as the Spearman correlation of the inferred LFCs between the two slides (estimated independently). All methods showed similar reproducibility. Focusing on *plausibility* scores, however, showed that both the *manual* benchmark and CSDE achieved higher values than the *automated* benchmark. The lower performance of the *automated* benchmark is likely a consequence of segmentation errors (resulting in misassignment) and classification errors (obscuring the DE signal by adding non T-cells).

For better interpretation of these results, we next analyzed the specific genes that were identified by each method (Figure 3d). The *Automated* benchmark produced several unexpected discoveries not detected by CSDE. These included genes characteristically expressed by stromal cells (*COL1A1*) and myeloid cells (*LYZ*), as well as other transcripts confirmed to be absent from T cells in this dataset (e.g., *CSF1, VEGFB, PECAM1*; Table S4).

Conversely, exploring the genes identified by the *manual* benchmark and CSDE revealed putative programs of T cell activation, exhaustion, and residency (Figure 3e). Both methods identified upregulation of key effector molecules (*GNLY, GZMB, PRF1*) indicating a potent cytotoxic capacity [30]. This was counterbalanced by signs of exhaustion, through tumor-enhanced expression of inhibitory receptors *LAG3* and *HAVCR2* (TIM-3), a well-described feature of exhausted T cells [31]. Concurrently, the downregulation of the lymph node-homing chemokine receptors *CCR7* and *CXCR4* suggested a shift towards a tissue-resident memory phenotype [32, 33].

CSDE uniquely identified additional gene expression changes that refined this characterization. It revealed a strong proliferative signal, marked by upregulation of the G1/S regulator *CCND1* and downregulation of the CDK inhibitor *CDKN1B* (p27) [34]. CSDE also identified upregulation of the anti-apoptotic factor *BCL2L1* (Bcl-xL) [35, 36], a key survival signal allowing T cells to persist in the tumor microenvironment by counteracting activation-induced cell death. This survival program is likely driven by the concurrent upregulation of transcriptional regulators NF-*κ*B (including the noncanonical component *NFKB2*) and *STAT3*, both known to drive *BCL2L1* expression [37, 38]. In contrast, downregulated *ZAP70* and *CD28*, consistent with impaired proximal TCR and costimulatory signaling in exhaustion, suggest chronic activation-associated attenuation of signaling capacity [39, 40, 41]. Finally, the downregulation of *GZMK* alongside upregulation of cytotoxic granzymes (*GZMH, GZMB*) is consistent with a shift toward a more cytotoxic effector program in intratumoral CD8 T cells [42, 43].

Collectively, these findings suggest that CSDE leads to biologically relevant discoveries while largely avoiding spurious findings likely resulting from automated artifacts.

#### 3.2 CSDE identifies hypoxic T cells in colon cancer

To strengthen the validity of our claims, we applied the same methodology to another cancer type, colon cancer, and for the same problem of characterizing intratumoral CD8 T cell phenotypes. On this problem, the automated pipeline produced a main dataset of 806,291 cells (Figure 4a, S3a). We followed the same procedure as for the lung data to annotate these segmented cells, identify tumor regions, and manually validate 600 cells randomly sampled from the slide (Figure S3b).

**Figure 4:**
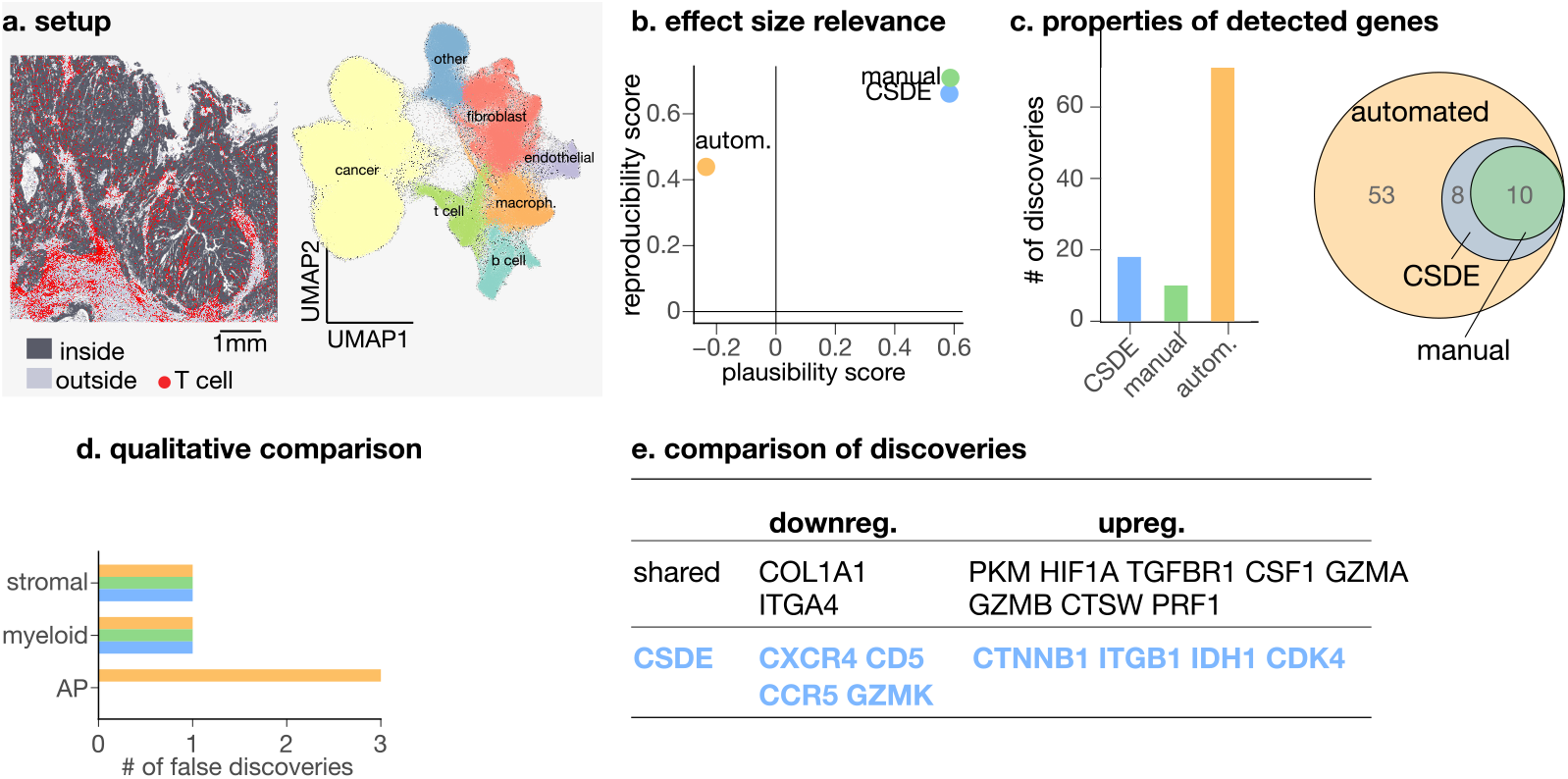
Colon experiment. Comparison of CD8 T cell gene expression profiles inside and outside tumors. **a**. *Left*: visualization of tumors in the tissue with CD8 T cells (obtained via automated pipeline) in red. *Right*: low-dimensional embeddings (UMAP) of the cells obtained by the automated pipeline. **b**. Relevance of the DE effect sizes obtained with the different methods based on reproducibiliy and plausibility scores. The biological plausibility score assesses the extent to which genes with large positive or negative LFCs are known to be expressed by T cells. The reproducibility score assesses the extent to which the LFC estimates are consistent between the two slides. See Supplementary Methods C.4 for more details. **c**. Properties of the discoveries made by the different methods. We compare the discoveries made by the different methods based on their number (*Left*) and overlap (*Right*). **d**. Quantification of non-T cell marker hits for the discoveries made by the different methods. AP: Antigen-presenting. The list of non-T cell marker genes is provided in Table S3. **e**. Comparison of the discoveries made by our method and the *manual* approach for the comparison of CD8 T cells inside and outside tumors.

### Comparative evaluation

Comparative benchmarking confirmed that CSDE provided more reliable estimates than *automated* and increased statistical power compared to *manual*. Focusing on comparing LFC estimates against a reference single-cell RNA-seq dataset ([29]; Supplementary Methods), CSDE again provided more plausible estimates than *automated* (Figure 4b). Comparing the number of detected DE genes, CSDE yielded an intermediate number of discoveries, almost twice as many as the *manual* approach but significantly fewer than *automated* (Figure 4c). Consistent with the lung cancer results, all genes detected by the *manual* method were also recovered by CSDE. As previously observed, *automated* detections were enriched for non-specific signals likely originating from the tissue microenvironment (Figure 4d). These included canonical stromal markers (*COL1A1, CSF1*), broad stress-response and metabolic genes (*SOD2, NFKBIA, JUN, JUNB*), as well as MHC class II genes (*HLA-DRA, HLA-DMA*). While some of these genes can be expressed by activated T cells, their broad enrichment in the *automated* discoveries suggests that the automated pipeline is particularly prone to contamination from ambient RNA and neighboring tumor or stromal cells. Notably, the few contaminants detected by CSDE (*COL1A1, CSF1*) were also present in the manual baseline, indicating that they likely represent irreducible background noise that cannot be corrected by the fast annotation protocol we employed in this work.

### Characterization of infiltrating T cells

Focusing on the discoveries made by the *manual* benchmark (all of which were recovered by CSDE), we identified a core tumor-infiltrating CD8 T cell profile characterized by cytotoxicity, metabolic adaptation to hypoxia, and immune evasion (Figure 4e). This shared profile included a cytotoxic state marked by *GZMB* and *PRF1*, consistent with prior characterizations of cytotoxic TILs in colorectal cancer [44, 31]. These CD8 T cells also showed features of metabolic stress and hypoxia, evidenced by the upregulation of *HIF1A* and *PKM. HIF1A* encodes the master regulator of the hypoxic response, which drives the expression of glycolytic enzymes including *PKM*. This coordinated upregulation suggests a metabolic adaptation enabling infiltrating CD8 T cells to maintain effector function and survival within the hypoxic, nutrient-restricted tumor microenvironment [45, 46, 47]. We also observed upregulation of *TGFBR1*, which encodes a receptor for the cytokine TGF-*β*. TGF-*β* is a central driver of immune evasion in colorectal cancer, promoting T cell exclusion and inhibiting their effector function [48, 49].

The specific discoveries made by CSDE described a richer characterization of the CD8 T cell population infiltrating the tumor. Refining the definition of the cytotoxic phenotype, the specific downregulation of *GZMK* alongside high *GZMB* indicated a shift toward a terminal effector state [44]. The downregulation of *CXCR4* and *CCR5* suggested a tissue-resident profile, consistent with the loss of migratory potential required for local tumor retention [50]. Crucially, CSDE also uncovered a distinct proliferative, stem-like state, marked by the upregulation in intratumoral CD8 T cells of *CDK4* and *CTNNB1. CDK4* encodes a key G1/S cell-cycle kinase driving proliferation, while *CTNNB1* encodes *β*-catenin, the central effector of Wnt signaling. The co-expression of these genes points to a Wnt-driven stem-like program, a state known to arrest terminal differentiation and sustain the proliferative capacity of tumor-infiltrating T cells [51, 52, 53].

## Discussion

This paper introduces CSDE, a framework for spatially-resolved differential expression that ensures statistical validity despite the segmentation and classification errors that are characteristic of automated pipelines. By integrating a large set of automated quantifications with a small, efficiently curated manual sample, CSDE corrects systematic biases without sacrificing statistical power. This approach empowers researchers to move beyond artifacts and focus on genuine biological signals. We demonstrate CSDE’s utility on lung and colorectal cancer datasets, where it revealed detailed profiles of intratumoral T cells, including specific signatures of hypoxia, exhaustion, and stem-like proliferation that were obscured by errors in the automated analysis.

CSDE uses a simple GLM to estimate LFCs, a choice that prioritizes interpretability and robustness. While this model effectively filters spurious discoveries, the framework could be naturally extended to incorporate richer noise models for overdispersion or to explicitly model spatial dependencies [54, 55]. However, the primary innovation of CSDE lies not in the base model itself, but in the rigorous integration of manual curation with automated quantifications to produce reliable estimates of DE.

CSDE minimizes annotation effort by validating automated outputs rather than generating annotations from scratch. Annotators inspect a small random sample to verify segmentation and correct labels. This strategy avoids manual segmentation, a slow process prone to errors that often fails to resolve ambiguous boundaries without transcriptomic context [19]. By validating fully-quantified cells instead, CSDE evaluates the entire preprocessing pipeline, capturing errors in both segmentation and classification. The rejection mechanism also serves as a quality control filter, identifying data issues such as ambient RNA, z-plane overlaps, or false-positive transcript calls [8].

While the statistical backbone of CSDE is available as a Python package ^2^, the implementation of a user interface for manual annotation poses additional challenges from a software engineering perspective, with issues relating to usability, stability of the interface, and its compatibility across input formats and coding environments. In this work, we relied on static diagnostic images loaded into CVAT [26], a standard tool for computer vision. While effective as a proof of concept, this approach has limitations: it lacks interactivity (e.g., exploring gene expression or z-planes) and requires custom scripts to handle technology-specific data formats. Emerging frameworks such as SpatialData [56] are establishing the necessary infrastructure to address these challenges by standardizing data access across platforms. As these ecosystems mature, they will enable the creation of technology and pipeline-agnostic interfaces for validation, ultimately democratizing human-in-the-loop quality control in spatial biology.

Finally, our view in this work is that expert time is better spent evaluating the quality of model predictions rather than attempting to replace the predictive components of the pipeline. This approach is not incompatible with efforts to improve the prediction models that compose preprocessing pipelines [57]; rather, it provides a safety net that ensures validity regardless of the underlying model’s performance. Looking ahead, a key direction is to make this limited annotation effort more efficient. For instance, rather than sampling cells randomly, future iterations could prioritize cells where the automated pipeline is uncertain, thereby targeting expert attention where it provides the most statistical leverage.

## Code availability

The source code for the `CSDE` Python package is available at https://github.com/YosefLab/CSDE. The code and notebooks required to reproduce the results and figures presented in this paper are available at https://github.com/PierreBoyeau/csde_reproducibility. This repository includes scripts for raw data download, segmentation, automated annotation, and the extraction of image crops for manual validation in CVAT, for the two datasets used in the paper.

## Supplementary Information

### A Supplementary Figures

**Figure S1:**
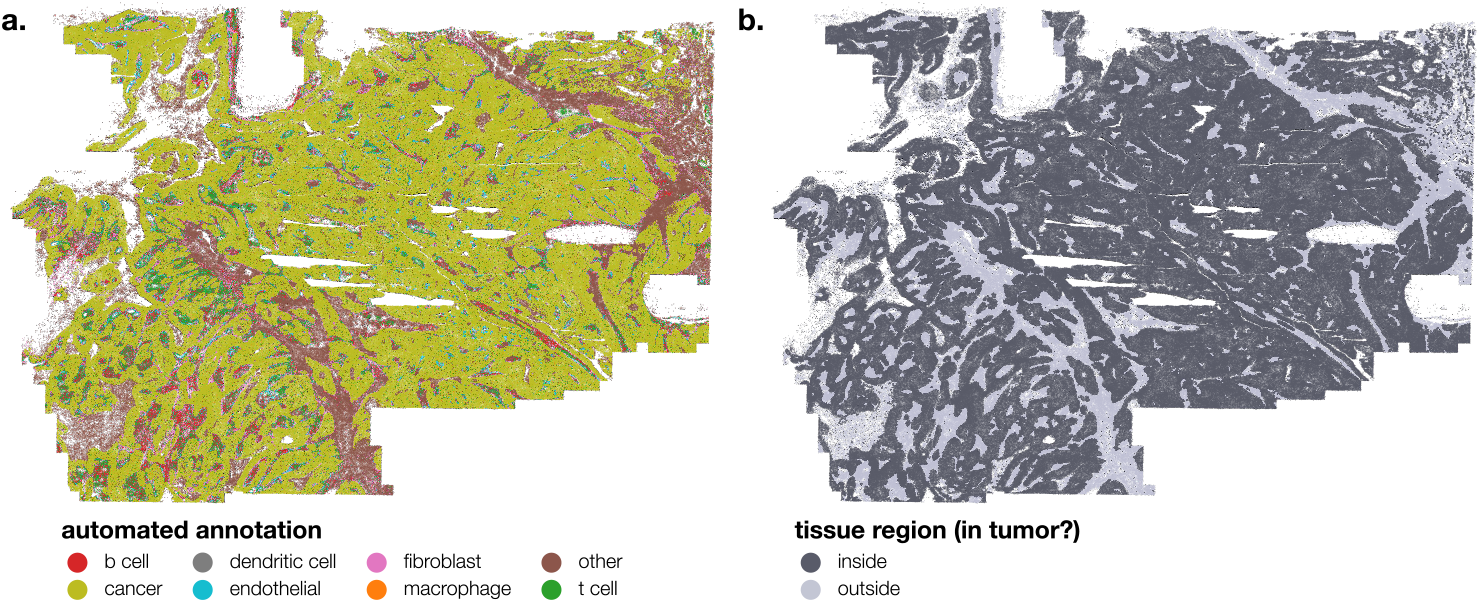
Visualization of cells in the main lung slide. colored by (**a**) ML-derived cell-type and (**b**) tumor region assignments.

**Figure S2:**
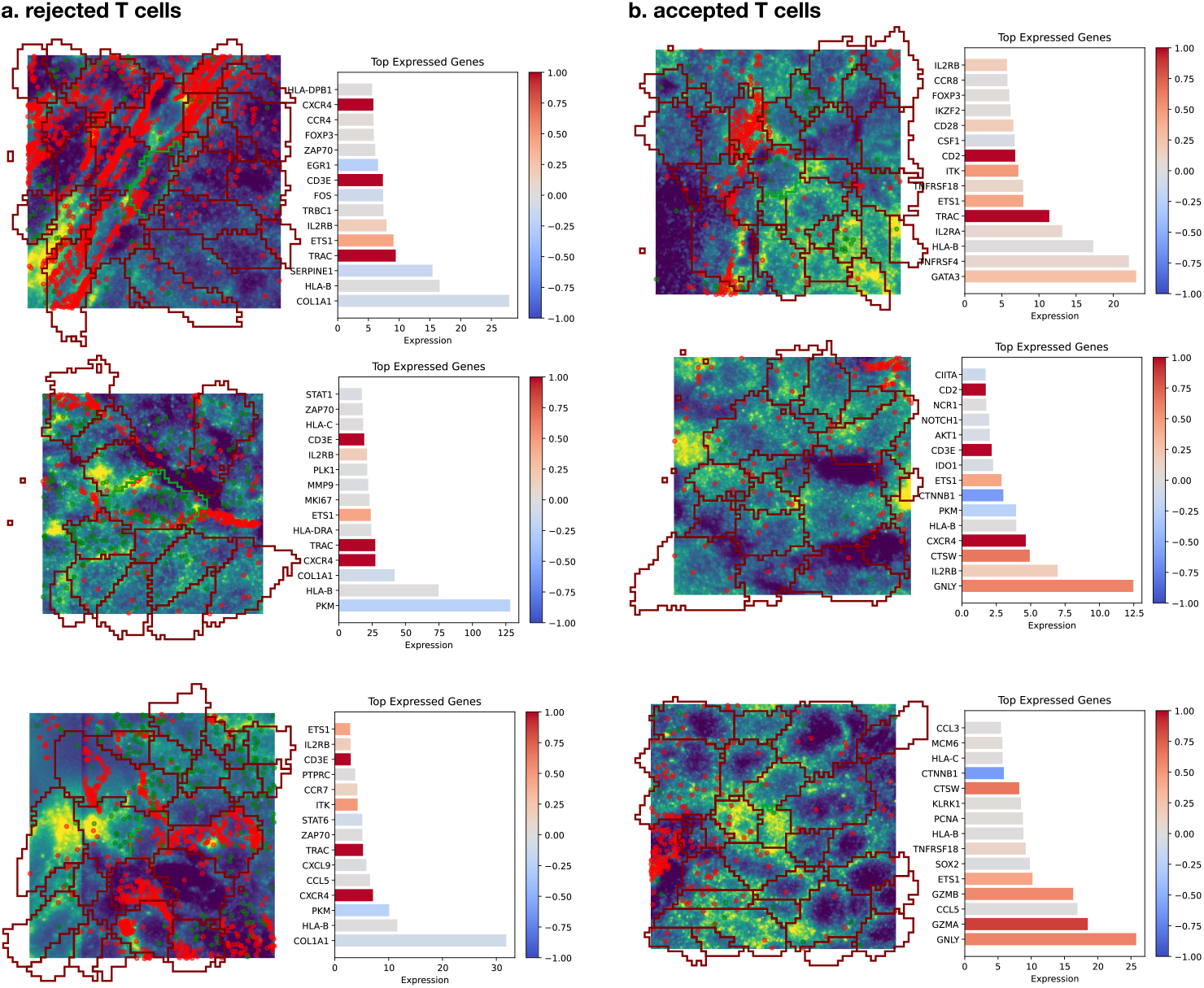
Cells with their manual annotations in the main lung slide. **a**. Cells annotated as T cells by the ML pipeline and rejected by the manual annotator. **b**. Cells annotated as T cells by the ML pipeline and accepted by the manual annotator. In both cases, we display three examples of cells from the same tissue slide. Each example shows cell nuclei (DAPI) along with the proposed ML segmentation overlay, as well as the top expressed genes in the cell.

**Figure S3:**
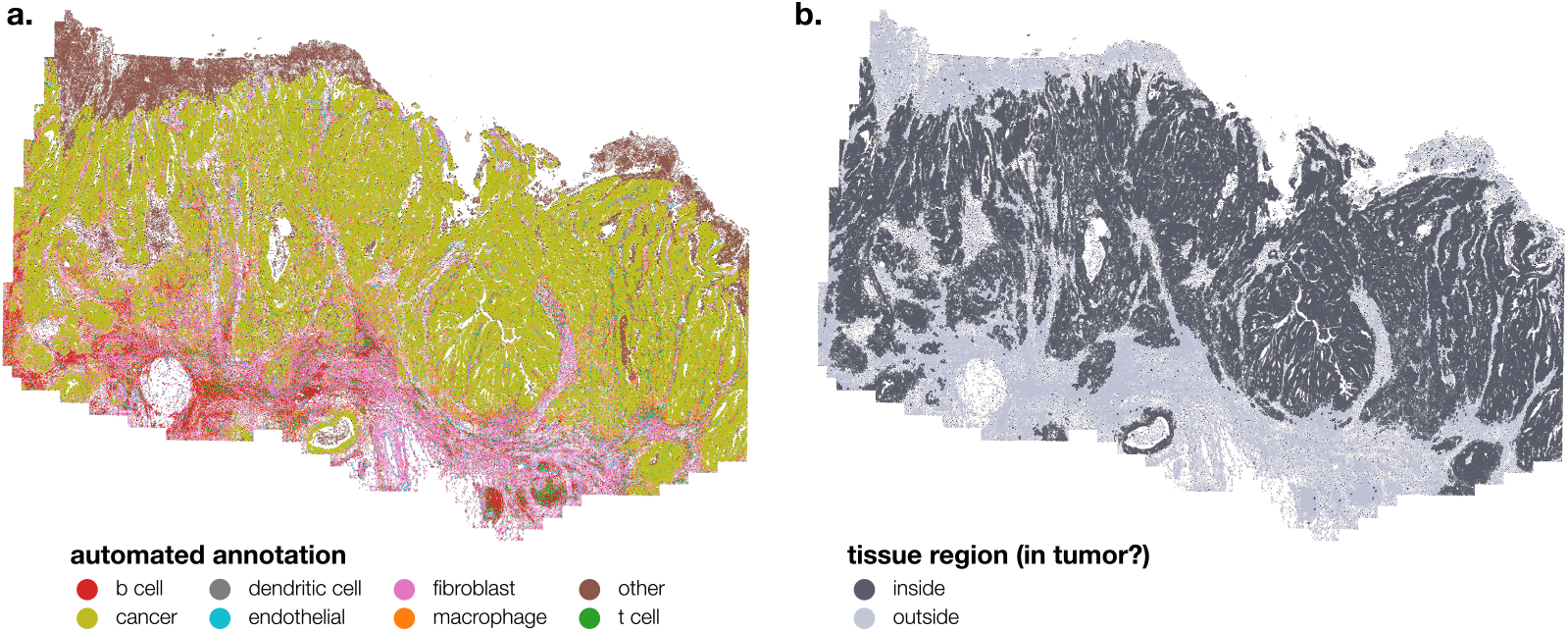
Visualization of cells in the main colon slide. colored by (**a**) ML-derived cell-type and (**b**) tumor region assignments.

**Figure S4:**
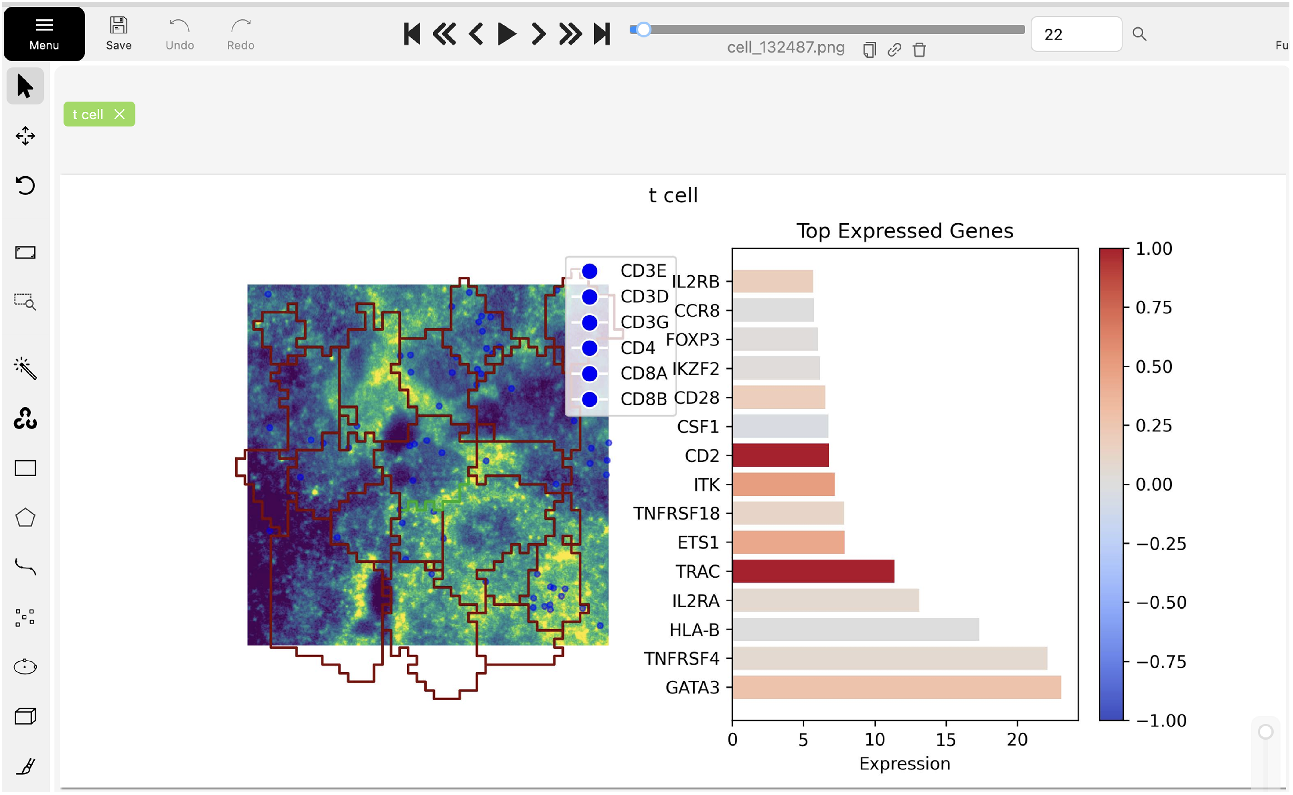
CVAT interface for manual annotation. The interface displays a representative cell panel (center), the assigned label (top left), and the queue of cells to be annotated (top). Since cell labels are predefined, the annotator can assign a label with a single key-press and navigate through the stack using arrow keys.

### B Supplementary Tables

**Table S1:** Markers employed to label clusters.

| Cell-type | Genes |
| --- | --- |
| Tumor | EPCAM, EGFR, VEGFA, HIF1A, MUC1, CEACAM1 |
| T cell | CD3E, CD3D, CD3G, CD4, CD8A, CD8B |
| NK cell | NCAM1, NKG7, KLRK1, GNLY, PRF1 |
| Dendritic cells | CD1C, CLEC4C, CD83, LAMP3, CD209 |
| Fibroblast | COL1A1, FAP, PDGFRA |
| Neutrophil | S100A9, FCGR3A, CXCR2, ELANE, MPO |
| Endothelial | PECAM1, CDH5, VWF, PLVAP |
| Macrophage | CD68, CD163, MRC1, MSR1 |
| B cell | CD19, CD79A, MS4A1 |

**Table S2:** Cell counts from ML-derived segmentation and classification. The table shows the total number of cells and the subset of T cells identified by our pipeline across four tissue slides.

| Slide | Total cells | T cells (%) |
| --- | --- | --- |
| Lung 1 | 351,309 | 19,867 (5.7%) |
| Lung 2 | 825,527 | 73,374 (8.9%) |
| Colon 1 | 671,075 | 5,390 (0.8%) |
| Colon 2 | 806,291 | 35,559 (4.4%) |

**Table S3:** List of non-T cell marker genes employed in the analysis.

| Category | Genes |
| --- | --- |
| Myeloid and DC | CD14, CD163, FCGR3A, MARCO, MRC1, TREM2, S100A9, LYZ, CYBB, CSF1, CSF1R, CD83, CD1C, CD1B, CD1E, LAMP3, CD207, CLEC4C, SIGLEC1, CD209 |
| Stromal | COL1A1, PECAM1, COL4A1, COL5A1, COL6A3, COL11A1, FN1, LAMB3, LAMC2, FBLN1, TNC, ACTA2, FAP, DES |
| Endothelial | VWF, CDH5, CLDN5, PLVAP, MMRN1, MMRN2, CLEC14A, KDR, TEK, VEGFB, VEGFA, VEGFC, ANGPT1, ANGPT2, NRP1 |
| Epithelial | EPCAM, CDH1, MUC1, MUC4 |
| Antigen presentation | HLA-DRA, HLA-DRB1, HLA-DPB1, HLA-DMA, HLA-DPA1, HLA-DQA1, CD276, LGALS9, PDCD1LG2, NCR1, NCR3, KLRG1, XBP1 |

**Table S4:** scRNA-seq statistics for ambiguous genes detected by automated in the lung dataset. The statistics are obtained from a pancancer scRNA-seq dataset [29].

| Gene | LFC (T cell vs. Rest) | Percent Expressed |
| --- | --- | --- |
| *LYZ* | -0.38 | 5.5% |
| *COL1A1* | 0.03 | 1.6% |
| *VEGFB* | -0.00 | 1.5% |
| *CSF1* | 0.03 | 3.2% |
| *PECAM1* | -0.02 | 3.1% |

### C Supplementary Methods

#### C.1 Proofs

In this section, we provide proofs for the main results of the paper. We start by proving the asymptotic normality of the CSDE estimator, which is a direct consequence of the results stated in Theorem 1 of [23].

*Proof of Corollary 1*. The log-likelihood of the working model of Equation (4) for a given dataset 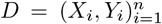, as a function of the optimization variable *b* (corresponding to parameter *β*), decomposes as:

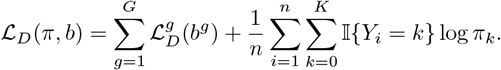

This decomposition shows that the MLE for *β*^*g*^ is obtained by maximizing 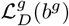 with respect to *b*^*g*^, independently of *π*. The CSDE estimator defined in Equation (5) maximizes the prediction-powered objective 𝒥_*g*_(*b*^*g*^), which is an unbiased estimator of the expected gene-specific log-likelihood 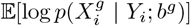. Since 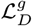 corresponds to a GLM, the objective 𝒥_*g*_(*b*^*g*^) is concave. We can therefore apply the results of [23] (which apply to convex M-estimators, or equivalently, maximum likelihood estimators for GLMs) to guarantee that the estimator is consistent and asymptotically normal.

We now prove the validity of the reweighting scheme used in Equation (7).

##### Proof of the validity of the reweighting scheme

Let 𝒫 denote the probability distribution over the population of cells in the tissue. Let 𝒬 denote the sampling distribution induced by the weighting function *w*, which is used to select cells for manual annotation. The importance sampling scheme described in the main text involves sampling a given observation *i*, with probability proportional to a user-defined weight *w*(ℐ _*i*_) based on the image ℐ_*i*_. Technically, the Radon-Nikodym derivative between the target and sampling distributions is proportional to the inverse of the weights

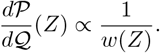

The reweighted objective function 𝒥_*g*_(*β*^*g*^) proposed in Equation (7) writes as

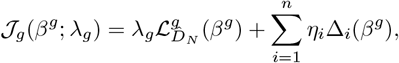

where 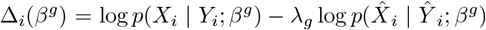 is the difference between the manual and weighted automated log-likelihoods for the *i*-th annotated cell. The weights are defined as 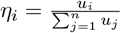, where *u*_*i*_ = 1*/w*(ℐ_*i*_) is the unnormalized weight of the *i*-th cell.

We analyze the two terms of the reweighted objective separately, focusing first on the imputation term 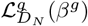. The unlabeled dataset 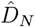 consists of *N* cells sampled uniformly from the population 𝒫. By the law of large numbers, as *N* → ∞:

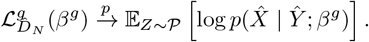

We now focus on the correction term 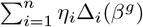. The manual dataset *D*_*n*_ consists of *n* cells drawn from *Q*, and the term 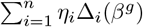 is a Self-Normalized Importance Sampling (SNIS) estimator. By the consistency of SNIS, as *n* → ∞, this term converges in probability to the expectation under the target distribution 𝒫:

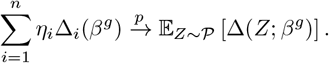

Combining these limits, the reweighted objective function converges to:

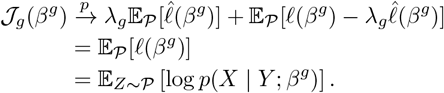

Thus, maximizing the reweighted objective 𝒥_*g*_ is asymptotically equivalent to maximizing the expected log-likelihood of the manual data on the full population, ensuring the consistency of the estimator 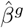.

#### C.2 Optimal selection of *λ*_*g*_

The hyperparameter *λ*_*g*_ ∈ [0, 1] controls the bias-variance trade-off by weighting the contribution of the automated dataset. Standard PPI approaches minimize the total variance of the parameter vector. However, since our primary interest is the log-fold change 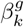 for a specific cell subset *k*, we select *λ* _*g*_ to minimize the asymptotic variance of this specific estimator, denoted as 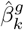.

To select *λ*_*g*_, we examine the asymptotic covariance matrix of the vector 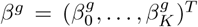, denoted as 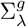. We first define the log-likelihood functions for the manual and automated data as

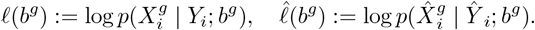

Following [23], the matrix 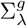 is given by:

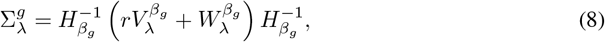

where 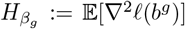 is the Hessian of the manual log-likelihood, and *r* is the ratio of the manual to automated dataset sizes. The terms 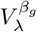 and 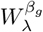 are defined as

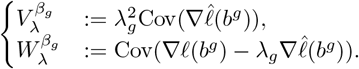

Note that the term 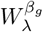 can be developed as:

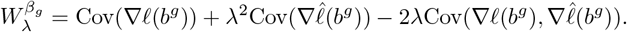

Our objective is to select *λ*_*g*_ to minimize the variance of the log-fold change for the subset of interest 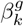, which corresponds to the (*k, k*)-th diagonal element of 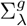, denoted as 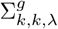. In other words, we look for *λ*_*g*_ minimizing the variance of 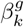:

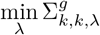

To minimize this quantity, we plug in the expressions for 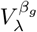 and 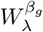, and reorganize terms in Equation (8) to obtain:

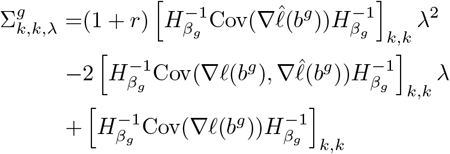

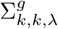 is quadratic with respect to *λ*, and the minimum has a closed-form solution given by:

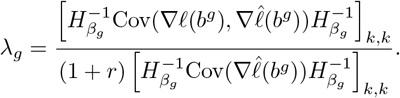

#### C.3 Annotation protocol

To ensure the quality and consistency of the manual dataset *D*_*n*_, we developed a standardized annotation protocol. The manual annotation workflow allows for the efficient validation of cell identity and segmentation. To maximize efficiency, we employ the importance sampling scheme described in the main text to prioritize the selection of potential cells of interest (here, CD8 T cells) for validation. However, the annotation protocol itself is applied uniformly to all selected cells. For each cell, we generate a composite image panel displaying nuclear staining (DAPI), relevant membrane markers, and key gene transcripts, overlaid with the automated segmentation boundaries (illustrated, for instance, in Figure S2). These panels are then uploaded to CVAT [26] for manual annotation, allowing the annotator to navigate through the cell stack rapidly (Supplementary Figure S4).

For each cell, the annotator makes one of three decisions:

- **Accept**: The automated segmentation is accurate, and the predicted label matches the cell’s expression profile and spatial context. The manual quantification is set to the automated one 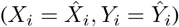.
- **Correct**: The segmentation is adequate, but the cell label is incorrect. The annotator assigns the correct label *c*, so *Y*_*i*_ = *c*, while retaining the automated expression 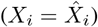.
- **Reject**: The segmentation is flawed (e.g., doublet, debris, poor boundaries). The cell is assigned to the “rejected” category (*Y*_*i*_ = *K*), effectively excluding it from the primary DE comparison.

This process, which requires a single key-press per cell, allows for rapid verification while accounting for the most common automated errors.

##### Implementation details

We implement the estimator of Equation (5) in a fast and parallelized manner using the `JAX` library, available at https://github.com/YosefLab/CSDE.

#### C.4 Benchmark

We benchmarked CSDE against two baseline approaches using four tissue slides from the MER-SCOPE platform (two lung cancer, two colon cancer) [25].

For these experiments, the automated pipeline utilized `proseg` [9] for cell segmentation and transcript quantification. Transcript counts were then normalized, log-transformed, and scaled to unit variance before applying PCA (50 components) and Leiden clustering [27] to identify cell clusters, which were then manually annotated at coarse resolution (Table S1). Clustering did not reliably distinguish between CD4 and CD8 T cells. To address this, we applied a consistent rule for all subsequent analyses: a cell annotated as a T cell (through either automated or manual methods) was further classified as a CD8 T cell if it expressed *CD8A* or *CD8B*, but not *CD4*.

Cells were stratified into tumor and adjacent regions based on their spatial proximity to tumor cells. Specifically, we first identified tumor cells using the automated annotations. We then computed the distance of each cell to its nearest tumor neighbor. Cells within a distance of 20*µm* were assigned to the tumor region, while the remaining cells were assigned to the adjacent region. The resulting tissue stratifications can be visualized in Figures S1 and S3. Crucially, these region assignments were computed once using the automated data and applied to all cells across all methods (Automated, Manual, CSDE). We therefore treat the cell locations and their spatial region assignments as fixed, focusing the correction on cell identity and gene expression.

We compared the following methods:

- **Automated**: A GLM (Equation 4) fitted solely on the large, automatically generated dataset. This represents standard practice but ignores segmentation/classification errors. We then inferred the asymptotic distributions of the GLM-based LFCs using a Wald test.
- **Manual**: A GLM fitted solely on the small, manually curated dataset (*n* ≈ 600 cells per slide), providing unbiased but high-variance estimates. Similar to the automated baseline, we inferred the asymptotic distributions of LFCs using a Wald test. This test is mathematically equivalent to PPI++ [23] with *λ* = 0, that is, discarding automated data and using only manual data for estimation.
- **CSDE**: Our proposed framework leveraging both datasets, as described in the main text.

Performance was evaluated using three metrics:

1. **DE Gene Identification**: The number of genes identified as differentially expressed with a p-value *<* 0.05 (Benjamini-Hochberg corrected).
2. **Biological Plausibility**: The Spearman correlation between the estimated LFCs and reference LFCs comparing CD8 T cells vs. non-T cells in a single-cell RNA-seq pancancer dataset [29].
3. **Reproducibility**: The Spearman correlation of LFC estimates for the comparison of interest (intratumoral vs. peritumoral CD8 T cells) between two replicate slides of the same tissue type.

#### Footnotes

1 For simplicity, we present these mappings as deterministic. However, our framework accommodates stochastic quantifications, where the output for a given raw data point is a random variable.

2 https://github.com/YosefLab/CSDE

